# Silk Fibroin as an Additive for Cell-Free Protein Synthesis

**DOI:** 10.1101/2020.05.28.121616

**Authors:** Marilyn S. Lee, Chia-Suei Hung, Daniel A. Phillips, Chelsea C. Buck, Maneesh K. Gupta, Matthew W. Lux

**Affiliations:** US Army Combat Capabilities Development Command Chemical and Biological Center, 8567 Ricketts Point Road, Aberdeen Proving Ground, MD 21010 USA; US Air Force Research Laboratory, 2179 12^th^ St., B652/R122 Wright-Patterson Air Force Base, OH 45433 USA; US Naval Research Laboratory Center for Bio/Molecular Science and Engineering, Bldg. 42, Room 303 4555 Overlook Ave. Washington, DC 20375; UES Inc., 4401 Dayton Xenia Rd., Beavercreek, OH 45432 USA

**Keywords:** Cell-free systems, Cell-free protein synthesis, CFPS, silk fibroin, preservation

## Abstract

Cell-free systems contain many proteins and metabolites required for complex functions such as transcription and translation or multi-step metabolic conversions. Research into expanding the delivery of these systems by drying or by embedding into other materials is enabling new applications in sensing, point-of-need manufacturing, and responsive materials. Meanwhile, silk fibroin from the silk worm, *Bombyx mori*, has received attention as a protective additive for dried enzyme formulations and as a material to build biocompatible hydrogels for controlled localization or delivery of biomolecular cargoes. In this work, we explore the effects of silk fibroin as an additive in cell-free protein synthesis (CFPS) reactions. Impacts of silk fibroin on CFPS activity and stability after drying, as well as the potential for incorporation of CFPS into hydrogels of crosslinked silk fibroin are assessed. We find that simple addition of silk fibroin increased productivity of the CFPS reactions by up to 42%, which we attribute to macromolecular crowding effects. However, we did not find evidence that silk fibroin provides a protective effects after drying as previously described for purified enzymes. Further, the enzymatic crosslinking transformations of silk fibroin typically used to form hydrogels are inhibited in the presence of the CFPS reaction mixture. Crosslinking attempts did not impact CFPS activity, but did yield localized protein aggregates rather than a hydrogel. We discuss the mechanisms at play in these results and how the silk fibroin-CFPS system might be improved for the design of cell-free devices.

## 1. Introduction

Cell-free protein synthesis (CFPS) is increasingly utilized as a platform to deliver genetic control over protein synthesis without the constraints of cell growth [1]. Avoiding transformation of DNA and culture growth time allows rapid prototyping of genetic circuits for design of sensors and enzyme cascades [2,3]. CFPS allows production of toxic proteins or molecules without the limits of cell viability [4]. Further, dried reaction components are able to be stored without refrigeration and rehydrated at the point of need to initiate activity [5–8]. These features allow the development of applications that function in austere environments without laboratory or cold-chain supply resources. One such application is eye-readable colorimetric sensing via CFPS reactions spotted onto paper, to sense many molecules from heavy metals to nucleic acid sequences in a fashion similar to pH paper [6,7,9–12]. Another application is on-demand drug production at the point of need [8,13–16]. In addition to dried formats, a few studies have explored delivery of CFPS via hydrogels, which could eventually lead to applications in therapeutics and responsive materials[17–20].

There are several limitations to the current state of the art in CFPS reactions. While significant shelf stability has been reported in the literature, there have been variable levels of success between different labs in achieving shelf-stable lyophilized reactions [5,21,22]. This is possibly attributable to variations in environmental storage conditions like humidity or seemingly small differences in CFPS reaction preparation. Thus, improvements in robustness are needed to manufacture reliable systems. Another limitation of cell-free reactions as sensors are delays in signal generation. Any improvements in reaction kinetics or signal amplification to produce a readable result in 15 minutes or less would allow cell-free reactions to be competitive with lateral flow-type field detection devices. Moreover, there has been limited investigation into incorporation of CFPS into other materials, such as hydrogels, to enable delivery of CFPS in new form factors like coatings, or to control localization of molecules in a reaction.

With these limitations in mind, supplementing excipients into the CFPS reaction can achieve improvements to activity and shelf life. Many different components and additives in CFPS reactions have been optimized to improve protein productivity. Some examples are magnesium and potassium salts, reducing and alkylating agents like DTT, glutathione, or iodoacetamide, or macromolecular crowding agents like polyethylene glycol, spermidine, or putrescine [23]. Macromolecular “crowders” impact transcription and protein folding processes thermodynamically and kinetically through changing viscosity and excluding volume to create environments similar to the interior of a cell [24]. Macromolecular crowding is particularly important for the correct function of native promoter systems *in vitro* [25–28], though too much crowding could slow down diffusion enough to limit protein synthesis [29,30]. For stabilization of dried CFPS components, a number of additives have been explored with success [5,21]. The search continues for new additives that can advance CFPS productivity, kinetics, activity lifetime, or shelf life. Further, an additive that forms a hydrogel could open the door to the design of localized or immobilized reactions [17,18,20].

One possible additive that has not been tested with CFPS up to now is silk fibroin (SF), a material depicted in Figure 1. SF protein isolated from *Bombyx mori* silk worm cocoons improves the stability of several purified enzymes such as glucose oxidase, horseradish peroxidase, lipase, or organophosphorus hydrolase when combined in solution or dried to form a film or sponge [31–34]. Though the stabilizing effects of silk on proteins have been demonstrated both in solution and in the solid state, the exact mechanisms of silk stabilization are not fully understood [34]. Stabilizing effects of SF in solution are likely similar to crowding or preferential exclusion effects observed for PEG or dextran excipients and possibly include hydrophobic shielding due to SF’s amphiphilic character. Stabilization in the dry form is attributed to either water-replacement hydrogen bond interactions between SF molecules and entrained proteins or restriction of molecular mobility within the glass matrix [33,34]. An additional property of SF is that enzymatic crosslinking of SF by horse radish peroxidase (HRP) or tyrosinase can convert an aqueous SF solution into a hydrogel while maintaining a mild reaction environment [35,36].

**Figure 1.**
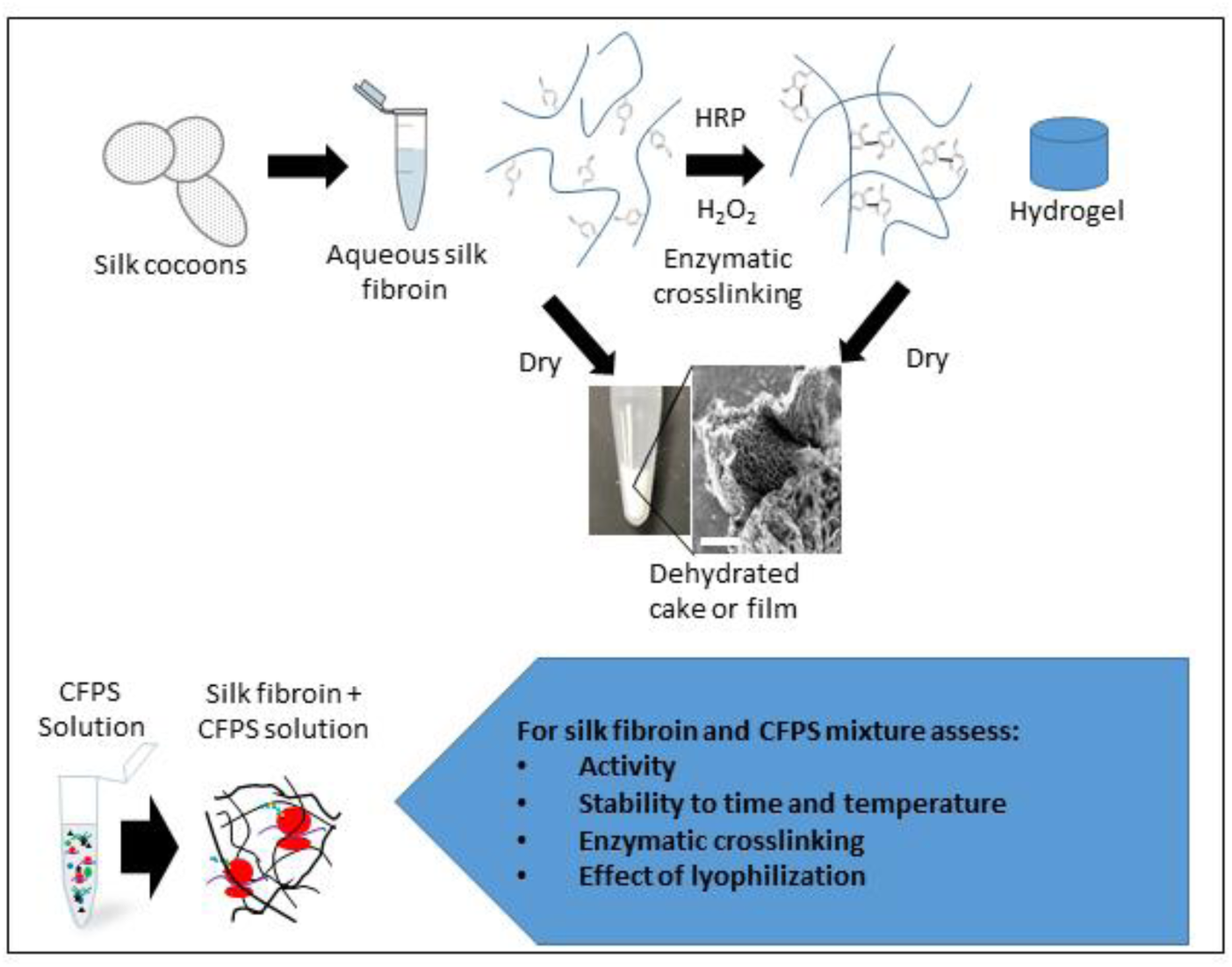
Flow chart representation of SF processing to achieve different materials. SF from silk worm cocoons is isolated and solubilized. Enzymatic crosslinking at tyrosine residues to forms a hydrogel. Either SF solution or hydrogel forms a porous film or sponge when dried. CFPS reaction components are entrapped in SF by mixing with the concentrated SF solution before crosslinking or drying.

In this study we sought to characterize the effects of SF on an *Escherichia coli* lysate-based CFPS system. Though there is evidence that SF can improve the function of individual purified proteins, these outcomes were previously uncharacterized for complex CFPS reactions. We anticipated three possible advantages to SF as an additive to CFPS reactions: increased activity through macromolecular crowding, increased stability after drying, and functionality within a hydrogel material. First, the effect of SF on CFPS activity in solution was assessed as a simple test for macromolecular crowding effects. Next, activity of CFPS reactions with added SF were assessed after freezing, lyophilization, or air-drying. Dried SF-CFPS films were challenged with long storage times and elevated temperatures to investigate any advantages for stabilization. In addition, the enzymatic crosslinking of SF into a hydrogel was examined in the presence of CFPS reaction components to determine whether SF-CFPS could be used to engineer localized reactions.

## 2. Materials and Methods

### 2.1 CFPS reaction components

Crude cell lysate for CFPS was derived from *E. coli* BL21 Star (DE3) as described previously [37]. An abbreviated method is provided here with a more detailed protocol available in the supplemental information. In short, bacteria were pre-cultured from frozen stock in 100 mL 2xYPTG media in a 500 mL shake flask at 37°C, 250 rpm for 16 hours. 5 mL of the pre-culture was used to seed 1 L of 2xYPTG media in a 2 L baffled shake flask that was shaken at 37°C, 250 rpm for approximately 3.25 hours, or until optical density at 600 nm reaches 3.0. Cells were then pelleted and washed with buffer. The wet weight of the pellet was then measured before the pellet was flash frozen in liquid nitrogen and stored at −80°C until further processing. For lysis, cell pellets were re-suspended in buffer and then lysed by sonication in 1.5 mL aliquots (Qsonica Q500, 20% amplitude, 40 seconds on, 59 seconds off, 540 J total energy). Lysates were then centrifuged and the supernatant was collected, aliquoted, flash frozen in liquid nitrogen, and stored at −80°C until reaction assembly.

The plasmid used for sfGFP expression in CFPS is pY71-sfGFP, which encodes super folder GFP under control of a T7 promoter and is selectable via kanamycin resistance [38]. The plasmid used to express Tyr1 tyrosinase from *Bacillus megaterium* via CFPS is pJV298-Tyr1 [39].

Expression from this plasmid is under the control of a *tac* promoter with a lac operator. Sequences for sfGFP and Tyr1 expression plasmids have been deposited on GenBank with identifiers MT346027 and MT346028, respectively. Additionally, plasmid maps are shown in Figure S1. Plasmid DNA is purified from transformed *E. coli* BL21 using a Promega PureYield plasmid midiprep kit followed by ethanol precipitation to further concentrate the DNA.

Unless otherwise noted, reagents were purchased from Millipore Sigma, St. Louis, MO. The final CFPS reactions contain the following: 12 mM magnesium glutamate, 130 mM potassium glutamate, 10 mM ammonium glutamate, 1.2 mM ATP, 0.85 mM each of GTP, CTP, and UTP, 34 ug/mL folinic acid, 170.6 ug/mL tRNAs, 2 mM of each of the 20 canonical amino acids, 33 mM phosphoenolpyruvate, 0.33 mM nicotinamide adenine dinucleotide, 0.27 mM coenzyme A, 1.5 mM spermidine, 1.0 mM putrescine, 4 mM oxalic acid, 57 mM HEPES, pH 7.4, 100 ug/mL T7 RNA polymerase, 6.5 nM plasmid DNA, 0.8 U/uL murine RNase inhibitor, and 26.7% (v/v) lysate. The “reagent mix” contains all of the above reagents apart from lysate and plasmid DNA. Aliquots of crude cell lysate are flash frozen in liquid nitrogen and stored at −80°C before use. Plasmid DNA, T7 RNA polymerase, and RNase inhibitor are stored separately at −20°C between uses. All other ingredients are pre-mixed, aliquoted, flash-frozen in liquid nitrogen, and stored at −80°C before use.

### 2.2 SF preparation

Regenerated SF is derived from the cocoon of the *Bombyx mori* silk worm (Mulberry Farms, Fallbrook, CA). The protocol for regenerated SF is adapted from Rockwood *et al*. [40]. The cocoons are delaminated into 2-3 layers, and finely sliced. Then, 7.5 g of delaminated cocoon are degummed by boiling for exactly 30 minutes in 3 L of 60 mM Na_2_CO_3_ solution to remove sericin and reduce the average SF molecular weight. Degummed SF fibers are removed and washed 3 times for 20 min in 2 L distilled water. SF is dried in a fume hood overnight and weighed to confirm removal of sericin, which is approximately 25% by weight of the initial cocoon material. The dried SF is then dissolved at a concentration of ∼20% (w/v) in 9.3 M LiBr. After dissolution, LiBr is removed by dialysis in water with a 12 kDa MWCO membrane. SF concentration is calculated from the dry weight of a known volume of solution. The solution is stored at 4°C until use with a shelf life of 1 month. SF degummed for 30 minutes has a molecular weight distribution centered around 100 kDa [40].

### 2.3 SF-CFPS reaction in microplate reader

Data in Figures 2 and 3 were collected from reactions performed at 25 µL total volume in 384-well microplate; all other reactions were performed at 15 µL volume. Lysate and reagents are stored frozen and thawed on ice before each experiment. For each reaction, lysate, reagent mix, DNA, RNase/DNase-free water, and SF stock solution (5% w/v stock concentration) are combined and thoroughly mixed by pipette. Final concentrations of each component can be found in Section 2.1 with the exception of SF, which is found in the corresponding figure or figure legend. When noted, HRP (Sigma Aldrich, p8375) and H_2_O_2_ are added to reach final concentrations of 80 U/mL and 0.005% (w/v), respectively. Crosslinking of SF solution alone occurs within a few minutes, so reactions were well mixed before addition of crosslinking reagents. CFPS reactions were performed in a BioTek Synergy H1 microplate reader with a monochromator light source maintained at 30°C with shaking for 8 hours. An adhesive film plate sealer (ThermoFisher 3501) was applied to cover wells and prevent evaporation. sfGFP fluorescence was monitored from the bottom of the plate with ex/em: 485/528 nm. To track SF dityrosine formation, fluorescence was monitored at ex/em: 300/425 nm. Relative fluorescent units were converted to µM sfGFP by measuring fluorescence of known concentrations of sfGFP (purified from *E. coli* by nickel column) on the same instrument and using the same measurement settings and plate conditions.

**Figure 2.**
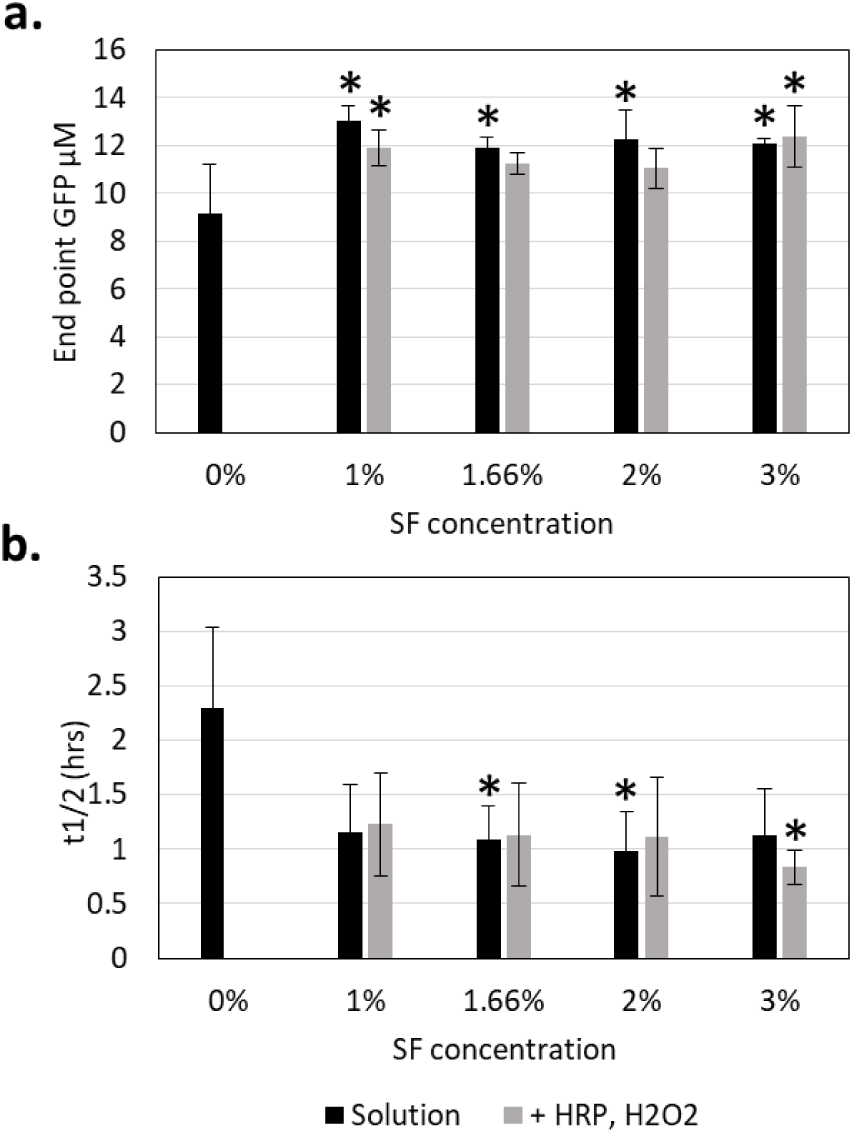
sfGFP production by CFPS in the presence of SF. (a) Comparison of relative sfGFP fluorescence at the endpoint of a CFPS reaction at 8 hrs. (b) Comparison of time to half maximal sfGFP fluorescence. Black bars are reactions where only SF is added. Grey bars are reactions containing SF as well as crosslinking reagents HRP and H_2_O_2_. Error bars represent the 95% confidence interval, n≥4. (* indicates p<0.05 compared to 0% (w/v) SF)

**Figure 3.**
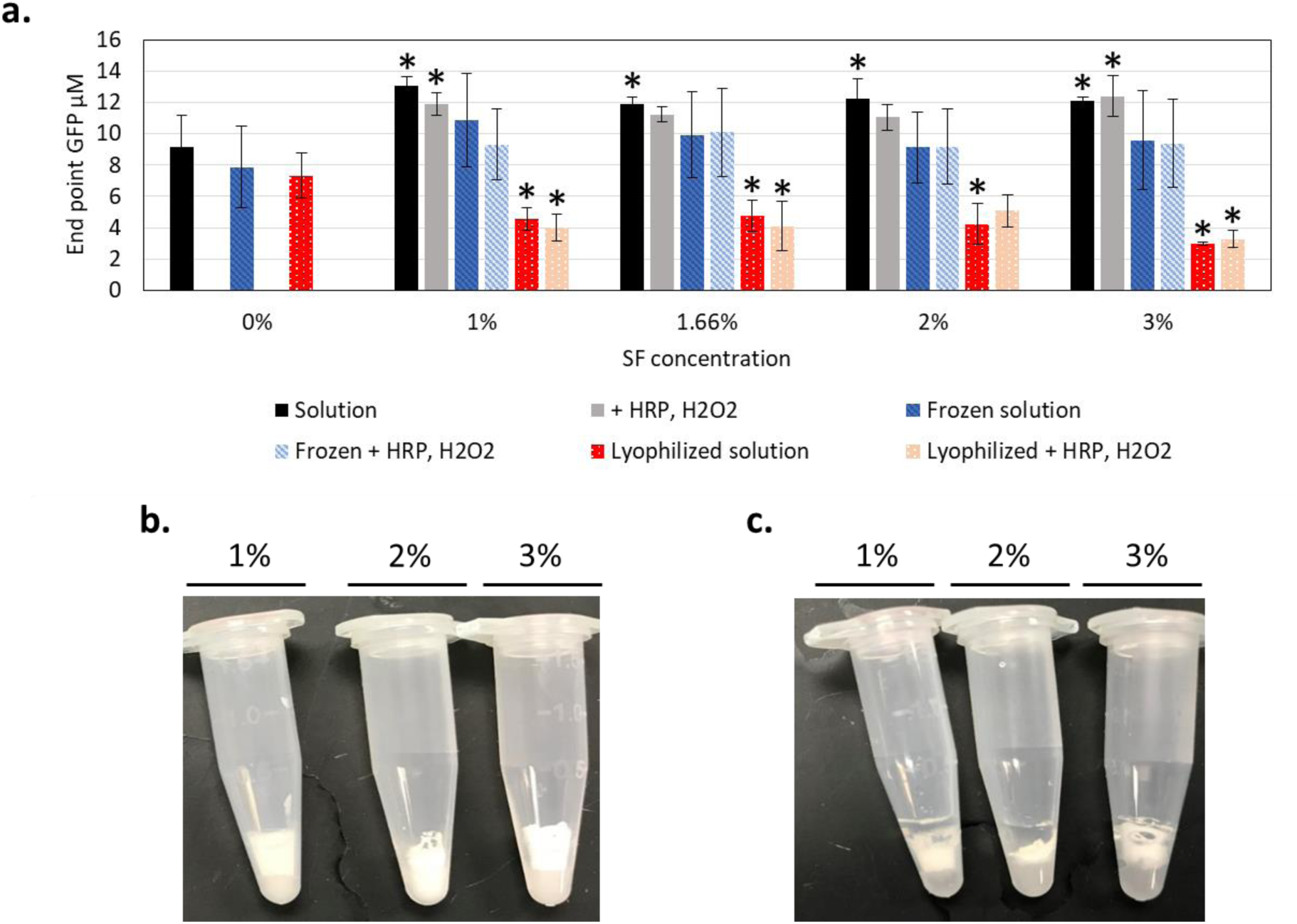
Freezing and lyophilization of CFPS reactions with SF solution (solution) or hydrogel (crosslinked). (a) Effects of freezing and lyophilization on CFPS activity measured by end point sfGFP fluorescence with and without presence of SF or HRP and H_2_O_2_. Error bars represent the 95% confidence interval, n≥4. (* indicates p<0.05 compared to 0% (w/v) SF) (b) Image of lyophilized SF fibroin crosslinked with HRP and H_2_O_2_ at various SF concentrations (c) Image of the samples in (b) after rehydration, total volume 100 µL.

### 2.4 Microscopy

To view the formation of sfGFP by CFPS embedded in the presence of SF, HRP, and H_2_O_2_, reactions were assembled as described previously. For sfGFP embedding, 18 µg of sfGFP (purified by nickel column) was crosslinked in 1.92% (w/v) SF before imaging. From the crosslinking reactions, 10 µL was quickly pipetted onto a 20 x 20 mm #1 cover glass (VWR, Radnor, PA, USA) and inverted onto a glass slide before the gelation reaction was completed. Acrylic nail polish was brushed around the edges of the cover glass to secure the coverslip to the slide. With one exception described below, confocal images were taken 4 hours post crosslinking on a Zeiss LSM 800 Airyscan confocal microscope with a Plan-Apochromat 40x/1.3 oil DIC (UV) VIS-IR M27 objective. sfGFP fluorescence was excited at 488 nm using 0.15% laser power, and the emission spectra was collected with 490-562 nm filters at 509 nm detected with an Airyscan detector. CFPS reactions and SF only controls were imaged with an 800-850V detector gain and a 2.0 digital gain. SF embedded sfGFP images utilized a 650 V detector gain and 1.0 digital gain to prevent oversaturated signals. Each image was taken with a 2.06 µs pixel dwell, 5.06 s scan time per frame. Images were collected of a 159.73 µm x 159.73 µm field of view with Zeiss Zen Blue imaging software (Carl Zeiss, LLC, Thornwood, NY, USA). The control sample containing CFPS reaction and crosslinking reagents HRP and H_2_O_2_ (no SF) was imaged using the same model microscope at a different location with some differences in equipment and settings. Images were collected using a Zeiss LSM 800 microscope with a Plan-Apochromat 20x/0.8 M27 objective in wide field imaging mode. The light source was a 475 nm Colibri 7 LED at 10% intensity, with filter set 90 HE LED. The image was captured with an Axiocam 506 camera and the exposure time was 150 ms. A representative frame was cropped from a 624.7×501.22 µm field of view.

### 2.5 Preservation by freezing, lyophilization, or air-drying

#### 2.5.1 Freezing

To preserve by freezing, all components are well mixed and distributed in microplate wells. Plates were kept on the bench for 10 minutes at room temperature to allow crosslinking reactions to proceed. Plates are then frozen at −80°C. Plates remained frozen for 5-7 days, then were thawed and incubated in a plate reader to measure GFP fluorescence as described in Section 2.3.

#### 2.5.2 Lyophilization

For lyophilized plates, reactions were first assembled and frozen at −80°C as described above. After one hour of freezing at −80°C, plates were transferred to a lyophilizer (Labconco cat#7948000) set to −40°C. The shelf temperature was then increased and maintained at −20°C for two hours, then raised again to 15°C for 17 hrs. The plates were then removed and were either rehydrated immediately or sealed and subjected to temperature challenges at room temperature, 37°C, or 50°C for the time points indicated. Reactions are rehydrated by adding RNase/DNase-free water equivalent to the total starting volume (15 or 25 µL), and measured with a plate reader as described in Section 2.3.

#### 2.5.3 Air-drying

For air-dried reactions, cell lysate and reagent mix were dried separately. Therefore, addition of SF to either the cell lysate or the reagent mix was tested in turn. Two controls with only water added to cell lysate or reagent mix were also constructed to differentiate the effect of adding SF from the effect of increasing the total volume before drying. Each reaction was 15 μL before drying. In the samples indicated, HRP and H_2_O_2_ were added as described in Section 2.3. The components were distributed into wells in a 384-well plate. Due to longer drying times at room temperature, plasmid DNA was not added before drying to prevent initiation of transcription. Plates were dried in a fume hood at room temperature for approximately 17 hours. After drying, plates were sealed and stored at room temperature, protected from light, for the indicated storage times. To initiate protein synthesis, components were rehydrated by adding water and plasmid DNA (total volume 15 µL) to the well containing the dried component (lysate or reagent mix) without SF, mixing, then transferring the mixture to the final reaction well containing dried component with SF. For example, in the case where reagent mix is dried alone and the cell lysate is dried with 1% (w/v) SF, the reagent mix is deposited in well “A”, while lysate, SF, HRP, H_2_O_2_, and water are deposited in well “B” before drying. To rehydrate, water and DNA is added to well “A” first, mixed, then transferred to well “B”. The sfGFP fluorescence of rehydrated and assembled reactions are monitored by plate reader as in Section 2.3. A “fresh” plate was constructed in parallel with the dried plates and run immediately without drying or storage time as a control.

### 2.6 Scanning electron microscopy

In 1.5 mL tubes, 100 µL of 1% (w/v) SF alone in solution and 100 µL of CFPS reaction containing 1% SF were incubated with HRP and H_2_O_2_ for 10 minutes at room temperature. The mixtures were then lyophilized as described above. The dried material in each tube was extracted with forceps and placed on carbon tape. The materials were imaged using a Phenom ProX desktop SEM.

### 2.7 Tyrosinase analysis

Tyrosinase from *Bacillus megaterium* was expressed via CFPS from the pJV298-Tyr1 plasmid template. 100 µM IPTG and 6.5 nM plasmid DNA are added to initiate tyrosinase production. To determine whether active tyrosinase is produced, activity was assayed for production of melanin from L-tyrosine. For this assay, 200 µM CuSO_4_ cofactor and 2 mM L-tyrosine substrate were added to the reaction for melanin formation. We found that adding 2 mM L-tyrosine at the start of the CFPS reaction incubation caused CFPS inhibition, so this ingredient was added after 2 hours of CFPS incubation to assay melanin formation. The SF crosslinking activity of CFPS-produced tyrosinase was then tested by adding 200 µM CuSO_4_ to Tyr1-CFPS reactions containing 1-3% (w/v) SF. To learn if diluting CFPS components would increase Tyr1 production of dityrosine crosslinks, 1, 5, or 10 µL of Tyr1-CFPS reaction was also diluted into SF solutions to reach 15µL total volume of either 1% or 3% (w/v) SF. The formation of dityrosine fluorescence was monitored at ex/em: 300/425 nm by microplate reader.

## 3. Results and Discussion

### 3.1 CFPS activity in the presence of SF

SF was added at 0-3% (w/v) final concentration to CFPS reaction mixtures containing cell lysate, resource reagent mix, and DNA. For some samples, crosslinking reagents (0.005% (w/v) H_2_O_2_ and 0.8 U/mL HRP) were added to test effects of these reagents on the CFPS reaction as a prelude to exploration of the potential formation of hydrogel materials (Section 3.3). Production of sfGFP from a plasmid template was measured over 8 hours (Figure 2a, Figure S2). Notably, addition of SF with and without HRP improved end point sfGFP signal, with activity increasing by 42% for 1% (w/v) SF without HRP. The result appears independent of the SF concentration across the 1-3% (w/v) range tested, though the result was not statistically significant for 1.66% or 2% SF (w/v) with HRP present. As an important metric for sensing applications, we also report the time to half maximal sfGFP signal where we similarly observed a decrease in the in the presence of silk, though fewer samples showed a statistically significant difference due to higher variability in this metric (Figure 2b).

Considering previous studies’ examination of the mechanism of silk stabilization, it is likely that this enhancement is due to volume exclusion of the SF molecules, an effect typical of macromolecular crowding [34]. The CFPS recipe used in this study includes previously optimized levels of spermidine and putrescine macromolecular crowding agents [41]. This recipe does not include PEG, which has been used for macromolecular crowding purposes in other studies. Further comparison between PEG and SF is needed to determine if they serve similar functions. In future work, studies of viscosity in the SF-CFPS system can shed light on the mechanism of enhancement, and measurement of mRNA levels could explore the hypothesis that the effect is related to enhanced transcription as observed in other studies. Importantly, this result establishes that SF can serve as a new type of beneficial additive for *in vitro* transcription and translation when maintained as a hydrated solution. Future studies could establish relative advantages and disadvantages of various additives for different CFPS recipes or applications.

### 3.2 SF-CFPS activity after freezing, drying, and storage

Flash freezing CFPS ingredients and storage at −80°C preserves activity indefinitely. However, a stringent cold chain is often not feasible for field applications. Room temperature storage of lyophilized or air-dried cell-free reactions maintains functional protein synthesis activity depending on the drying technique and CFPS recipe [5,6,13,21]. It is important to evaluate how SF may impact each of these preservation methods. CFPS reagents combined with SF were first subjected to freezing or lyophilization. The production of sfGFP from these reagents are shown in Figure 3a. A comparison is made between reactions with different SF concentrations and with or without HRP and H_2_O_2_. Reactions frozen with SF behaved similarly to samples without SF (p>0.05). In contrast, lyophilized reactions containing SF produced significantly less sfGFP than controls without SF. Drying SF by lyophilization resulted in an opaque white sponge material, as seen in other studies [42]. Rehydration of the mixture to initiate CFPS does not completely return the SF to its original soluble state; rather, much of the insoluble sponge remains, perhaps reducing the availability of resources trapped in the material (Figure 3b, c). Increasing concentrations of SF further decreased activity after lyophilization. The addition of HRP and H_2_O_2_ again did not affect the resulting sfGFP production across all of the conditions tested.

Next, lyophilized CFPS reactions were subjected to varying degrees of temperature stress to learn if SF provided improved stability to the dried CFPS reactions, despite an overall loss of activity. Samples with and without SF were stored sealed at room temperature (RT), 37°C, or 50°C for up to 72 hours. At each time point, samples were removed and rehydrated. sfGFP production after 8 hours at 30°C was measured via plate reader (Figure 4 a-c). The results confirm that activity of lyophilized material is relatively stable at room temperature over this time period, but decreases steadily at elevated temperatures. In contrast to results reported for many purified enzymes, in our tests the addition of SF did not significantly improve the stability of lysate-based CFPS reactions when lyophilized. It is important to note that samples were stored under atmosphere and without desiccant for these challenge experiments. Storing under inert gas or with desiccant may improve shelf life. It is possible to dry SF in such a way that the films are completely soluble when rehydrated [34], indicating that recovery and stabilization of SF-CFPS activity might be improved via optimization of the lyophilization procedure. Another consideration in stabilization is the ratio of SF to other proteins. We used concentrations of silk that were practical to achieve in solution without disrupting the concentrations of the other CFPS components. We estimate the ratio of lysate protein to SF protein in this experiment was on the order of 0.8 µg/µg, which is 1-3 orders of magnitude greater loading of protein cargo than is described in some previous studies on protein stabilization in soluble silk films [31,43]. Thus, reducing the ratio of cargo protein to SF could improve stabilization of the SF-CFPS activity.

**Figure 4.**
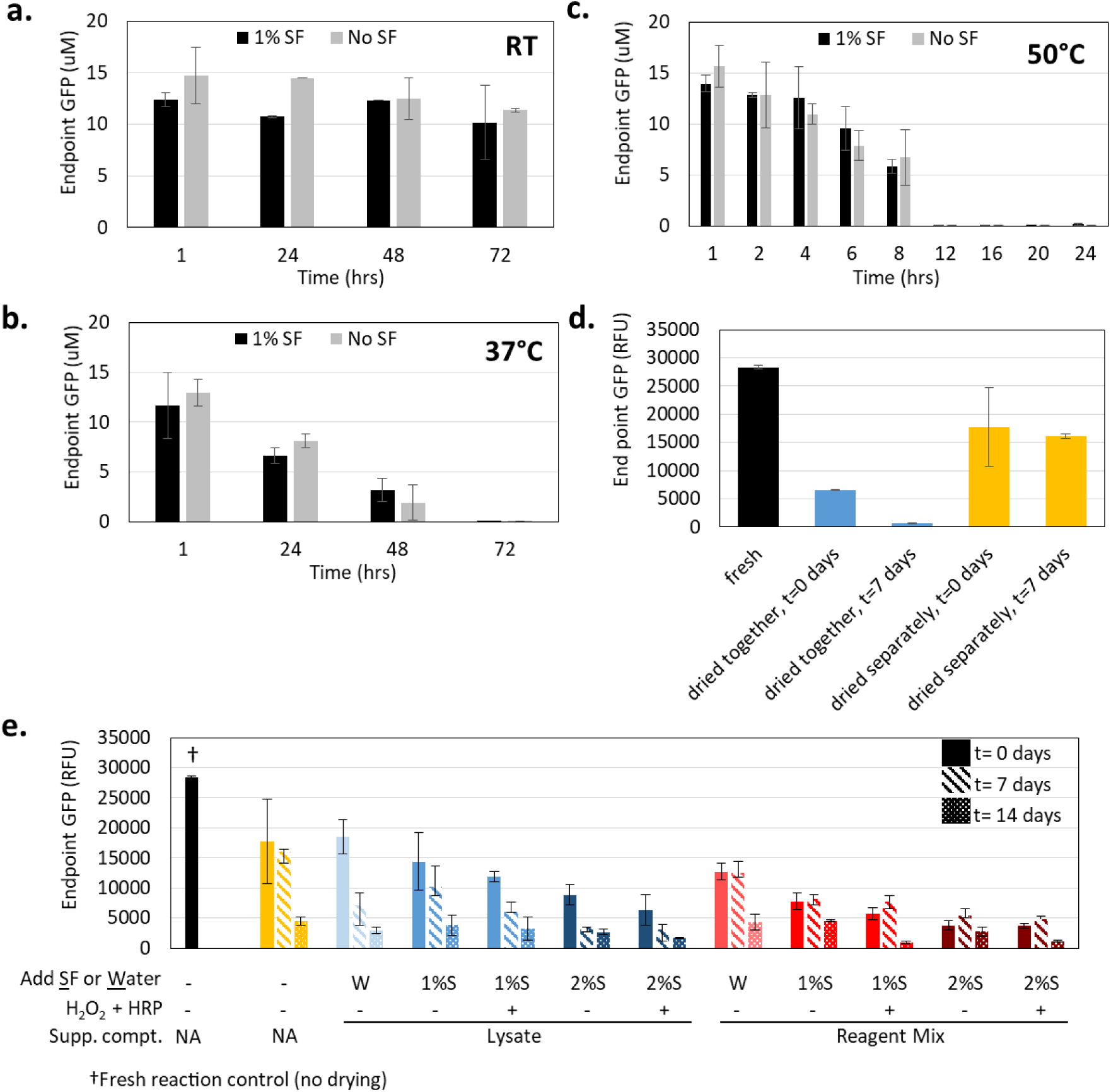
Stability of lyophilized SF-CFPS reactions to time and temperature. (a-c) Mean endpoint sfGFP levels of CFPS reactions containing 0% (grey) or 1% (w/v) (black) SF lyophilized, exposed to one of three temperatures (a=room temperature, b=37°C, c=50°C), and rehydrated. n≥2. d) Comparison of drying configurations for air-dried CFPS reactions. Reagent mix and lysate were either dried together or separately. e) Effects of air-drying and extended storage times on CFPS activity in the presence of SF solution or SF with HRP and H_2_O_2_. For each reaction the lysate and reagent mix were dried separately. The first row of labels shows whether water (W) or SF (S) was added before drying with a percentage to indicate the final concentration of SF. The supplemented component (Supp. compt.) refers to the component to which SF or water was added, either the lysate or the reagent mix. Error bars represent the 95% confidence interval, n≥3.

Air-drying is another method of storage that has the advantage of not requiring specialized equipment. Previous studies by Karig *et al*. demonstrated that CFPS reactions are best preserved if the lysate is dried separately from the other reaction components, though the formulation of CFPS reagents used in that study was different than the formulation used here [5]. We first confirmed in the absence of SF that drying cell lysate and reagent mix separately improved CFPS activity over drying together for our CFPS formulation (Figure 4d). Notably, this experiment showed little or no drop in activity after storage for 7 days, suggesting a significant improvement in stability; however, stability did deteriorate at 14 days in subsequent longer experiments (Figure 4e). We next assessed the effect of SF addition to either lysate or reagent mix before drying each separately at room temperature. Again, HRP and H_2_O_2_ were added to some samples containing SF. Because the added SF effectively diluted sample concentrations prior to drying, water was added to some samples as a control. Unlike in other experiments, DNA was withheld from these reactions prior to preservation due to the potential for reactions to initiate during the longer drying period. Function of the dried components was tracked for samples stored sealed at room temperature for 0, 7, or 14 days, and compared to fresh components that were not dried. To test activity, the separately dried lysate and reagent mixes were rehydrated and combined along with fresh DNA to initiate the formation of sfGFP. Air-drying under these conditions leads to a 37% drop in activity compared to a fresh reaction. Increasing concentration of SF in either the lysate or the reagent mix resulted in further decreased activity after drying, although the effect at 1% (w/v) SF concentration is minor (Figure 4e). Reactions with HRP and H_2_O_2_ added had similar or slightly less productivity than samples without. Looking at longer storage times, the stability of the dried reactions was not improved by addition of SF. In all cases, sfGFP production after 14 days of storage was reduced to about 18% or less of the fresh reaction. As with the lyophilized samples, we point out that other CFPS recipes, drying conditions, or additives could yield better stability both with and without SF [5]. In all, we conclude that SF does not improve stabilization of CFPS for the conditions we tested, but may still have that benefit in other situations.

### 3.3 Assessing SF crosslinking and hydrogel formation

HRP has been reported to catalyze the formation of dityrosine linkages between residues of SF to form a hydrogel from aqueous SF solution [44]. In this reaction, the iron-chelating porphyrin coordinated within the HRP active site is oxidized by H_2_O_2_, allowing the enzyme to accept electrons from phenol and aniline derivatives to form radicals. These radicals can then react to bridge surface exposed tyrosine residues of the SF and likely other proteins in the mixture [35,45]. For these concentrations of SF, H_2_O_2_, and HRP, crosslinking and hydrogel formation should occur in about two minutes or less. This was confirmed for pure SF solutions by vial inversion test (Figure S3). Curiously, the presence of CFPS reagents, or even just the lysate component, prevented hydrogel formation and caused failure of the vial inversion test. Dityrosine fluorescence measurement shed light on the progression of tyrosine crosslinking under different conditions (Figure 5) [35]. Addition of crosslinking reagents HRP and H_2_O_2_ to aqueous SF solution with no additional ingredients demonstrated a clear and rapid increase in dityrosine fluorescence. However, when CFPS components were included in the solution, no dityrosine formation was observed (Figure 5a). This indicated that the HRP enzyme activity on tyrosine was inhibited in the presence of CFPS reagents, either through some inhibitor molecule or competition with other substrates in the mixture. HRP has little substrate specificity, so it is quite likely that unintended reactions proceed in solutions with high complexity even though SF protein is a major component. It is also possible that the redox state of the reaction environment is not conducive to HRP activity. However, increasing the levels of HRP and H_2_O_2_ in the reaction up to ten times the original concentration did not result in any observable tyrosine crosslinking in the samples where CFPS components are included (Figure 5b). In fact, increasing the levels of HRP and H_2_O_2_ decreased dityrosine formation in solutions with SF alone, likely due to excess oxidation and inactivation of the HRP enzyme.

**Figure 5.**
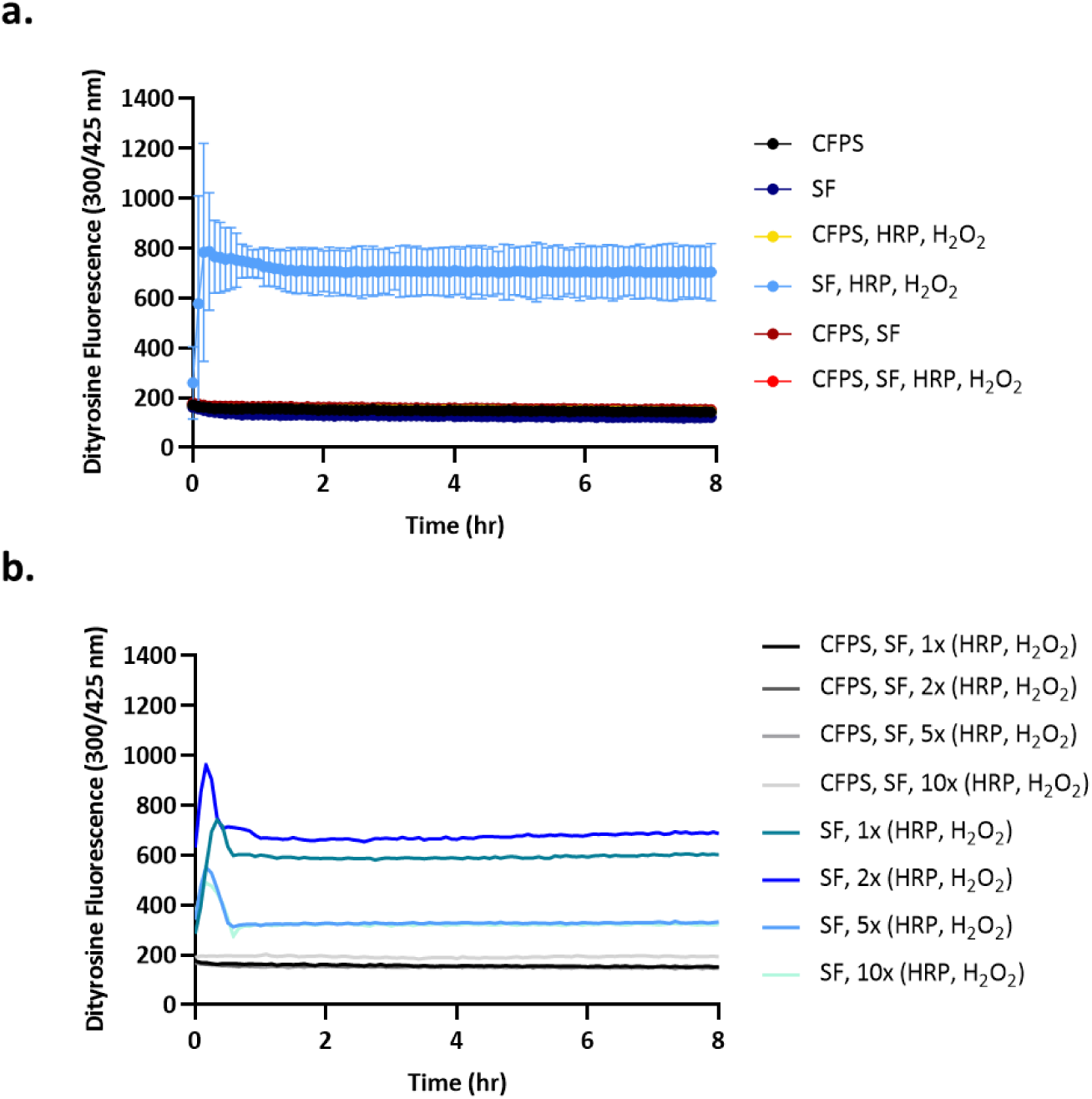
Microplate reader assay to detect appearance of dityrosine fluorescence. (a) Samples with or without CFPS, 1% (w/v) SF, or HRP and H_2_O_2_. Error bars represent the 95% confidence interval. (b) Samples with SF, with or without CFPS, and with increasing levels of crosslinking reagents HRP and H_2_O_2_.

Tyrosinase is an alternative enzyme that catalyzes dityrosine crosslink formation via the intermediates *o*-diphenol and *o*-quinone followed by radical formation and dityrosine bonding [36,45]. Tyrosinase has a copper center in the active site and catalyzes a different reaction than HRP. Its activity is also much more specific to mono- and di-phenols such as tyrosine [46]. To quickly learn if tyrosinase was active in a CFPS reaction mixture, plasmid DNA encoding tyrosinase from *Bacillus megaterium* was added to the SF-CFPS solution to initiate cell-free expression of tyrosinase enzyme. The production of active tyrosinase via CFPS was confirmed by the observation of melanin formation upon addition of CuSO_4_ and L-tyrosine (Figure S4). Nonetheless, no samples exhibited a detectable dityrosine signal when SF and CuSO_4_ were combined in a tyrosinase CFPS reaction, and the solutions remained viscous rather than forming gels. Because we used tyrosinase expressed in CFPS reactions rather than purified enzyme, we were not able to assess titration of the enzyme independent of the CFPS components as we did for HRP; however, titrating tyrosinase CFPS reaction into SF solution to dilute CFPS components did not improve the result. Like HRP, tyrosinase failed to crosslink SF in the presence of CFPS components.

In summary, these experiments demonstrate the detrimental effects of lysate and CFPS reaction mixtures on two different enzymatic techniques to modify SF substrate to form a hydrogel. Since the activity of tyrosinase was confirmed in the presence of CFPS components via melanin production there is evidence that competing side reactions may be the cause. Choosing a more targeted enzymatic crosslinking method could be critical to successful implementation in the complex CFPS reaction environment. Other factors that could influence HRP and tyrosinase activity may be tweaked in a cell-free system, including ionic strength or redox–related additives or enzymes such as reductases and dismutases [23]. The difficulty of reformulating the CFPS environment for each target enzyme is a significant hurdle to potentiating some sensitive enzymatic outputs via CFPS. As with the potential for SF to provide stabilization to dried CFPS reactions (Section 3.2), we conclude that SF did not offer a path to CFPS incorporation in a hydrogel for the conditions we tested, but further investigations might unlock this possibility.

### 3.4 SF-CFPS morphology

Next, we examined the morphology of the SF-CFPS reactions via observing sfGFP fluorescence in solution by fluorescence microscopy as well as imaging the dried cake material by SEM (Figure 6). Notably, when SF, HRP, and H_2_O_2_ are present sfGFP signal is isolated to ∼10-50 μm inclusions surrounded by regions with comparatively little fluorescence (Figure 6a). In contrast, if the SF is mixed with a CFPS reaction without crosslinking reagents (Figure 6f) or if crosslinking reagents are mixed with a CFPS reaction without SF (Figure 6b), sfGFP signal is uniform across the solution. Also, if SF is mixed with purified sfGFP and crosslinked with HRP and H_2_O_2_, the sfGFP remains uniformly distributed (Figure 6d) as does CFPS reaction alone without SF or crosslinking reagents (Figure 6c). This evidence indicates that, though HRP and H_2_O_2_ do not detectably catalyze tyrosine crosslinking and hydrogel formation, they are not inert in the presence of both CFPS reagents and SF.

**Figure 6.**
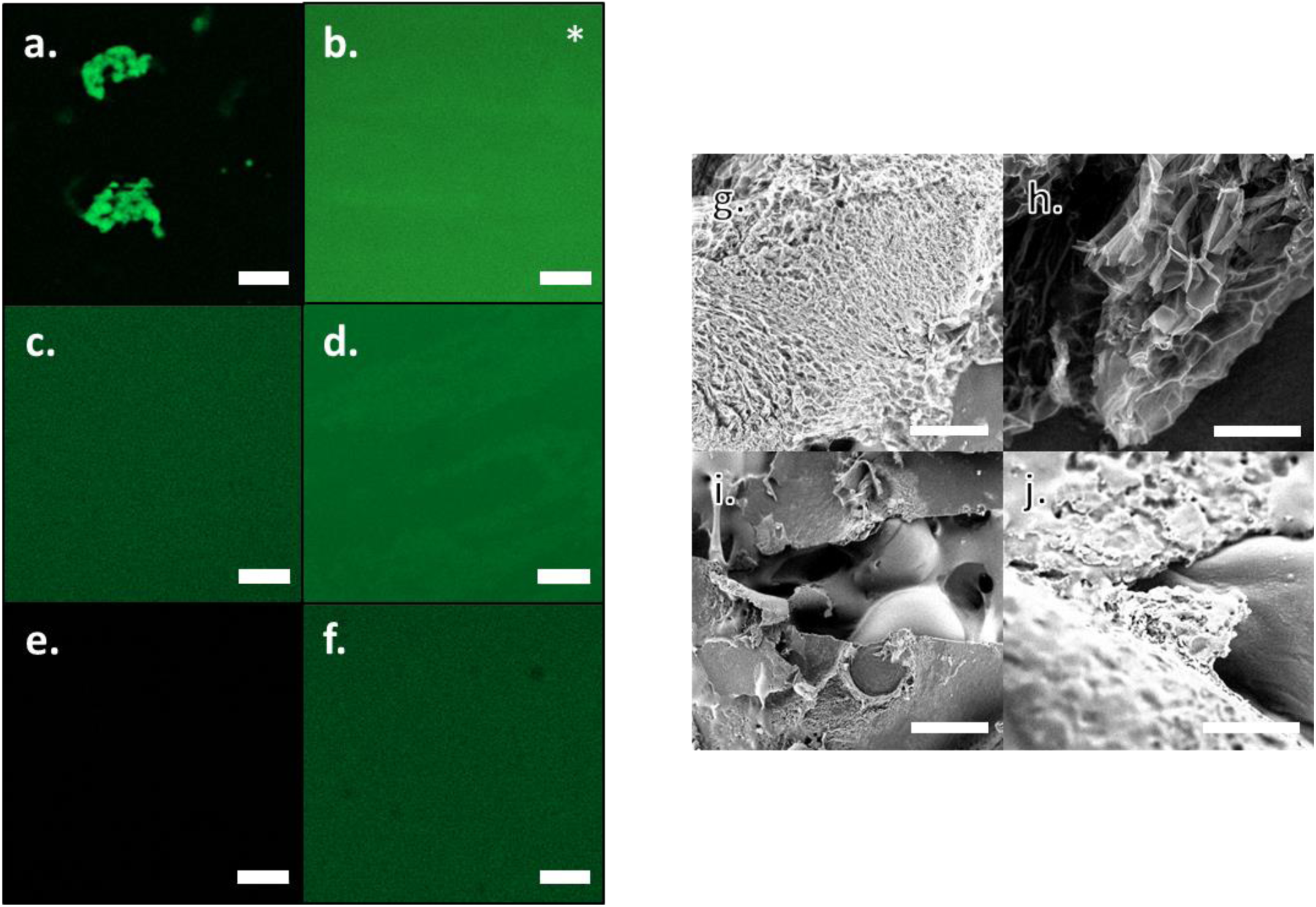
SF-CFPS morphology imaged via fluorescent microscopy and SEM. (a-f) sfGFP fluorescence of hydrated samples (scale bar 20 µm): (a) CFPS reaction containing 1% (w/v) SF, HRP, and H_2_O_2_; (b) CFPS reaction containing HRP and H_2_O_2_; (c) CFPS reaction; (d) purified sfGFP mixed with 1% SF, HRP, and H_2_O_2_; (e) background fluorescence with 1% SF, HRP, and H_2_O_2_; (f) CFPS reaction, 1% (w/v) SF, no crosslinking reagents. Images in panels (a), (c), (e), and (f) were collected at 800-850 V detector gain and 2.0 digital gain while panel (d) was collected at 650 V detector gain and 1.0 digital gain. Panel (b) has an asterisk because the image was collected using a different microscope as described in methods. (g-j) SEM images of lyophilized samples: (g, h) 1% (w/v) SF, HRP, H_2_O_2_ (f=scale bar 200 µm, g= scale bar 50 µm); (i, j) CFPS reaction, 1% (w/v) SF, HRP, H_2_O_2_ (h=scale bar 200 µm, i= scale bar 50 µm).

One possibility is that the inclusions do result from HRP-mediated tyrosine crosslinking of the SF, but that the presence of CFPS reduces the HRP activity on the SF to a level below the detection limit of the dityrosine fluorescence assay and vial inversion test. Alternatively, though not mutually exclusively, the inclusions could indicate that HRP is acting on substrates other than the SF protein. However, if so, the non-specific activity does not change overall sfGFP productivity with or without SF or cause enough aggregation in the absence of SF to be observed by our fluorescent microscope. It is also notable that phase separation of GFP from SF proteins has been previously reported in drop cast films from aqueous mixtures of SF and GFP [47]. Incomplete or non-specific crosslinking could easily disrupt SF solubility. The observed aggregates could also be some synergistic combination of effects, such as non-specific crosslinking of CFPS or sfGFP paired with macromolecular crowding effects of the SF to form inclusion bodies of sfGFP. We note that this interesting phenomenon introduces the possibility of using enzymatic crosslinking to create localization of functions in the cell-free environment. Previous studies have made use of hydrogels or molecular scaffolding to improve activity or organize functions into compartments, introducing diffusion controlled dynamics [48]. Titrating different concentrations of HRP did not improve the formation of a gel and was not explored here using microscopy, but could be a parameter useful for altering the degree of aggregation. The use of HRP crosslinking for this purpose merits future study.

SEM of lyophilized SF with or without CFPS components demonstrates how addition of CFPS components change the morphology of the dried material. SF hydrogel crosslinked with HRP and H_2_O_2_ forms a highly porous sponge material when lyophilized (Figure 6 f, g). On the other hand, when CFPS reagents are added in addition to SF and crosslinking reagents, the lyophilized cake is collapsed and non-porous (Figure 6 h,i). This observation yet again demonstrates that HRP activity on SF is different in the presence of CFPS. The change in structure could offer clues to potential future investigations to improve CFPS incorporation into hydrogel materials.

## 4. Conclusions

Motivated by expanding applications for CFPS [1], there is an ongoing push to improve the productivity, kinetics, and shelf life of CFPS systems. It is also important to identify the factors that control these properties and expand understanding of the underlying mechanisms. In this study, the putative advantages of SF fibroin for *in vitro* enzyme systems were tested for lysate-based CFPS reactions. The addition of SF to CFPS reactions in solution improves both protein productivity and kinetics. Volume exclusion and shielding properties are hypothesized benefits of SF that may explain the observed activity increase as a macromolecular crowding effect. As an unstructured protein excipient with unique properties, the effects of SF may be different than other additives such as PEG or sugar-based molecular crowding agents. In future work, mRNA and viscosity measurements on the SF-CFPS system may help elucidate this effect further.

On the other hand, SF did not improve the performance of dried CFPS reactions in our tests. SF-CFPS is tolerant to freezing, but has decreased activity after lyophilization or air drying. SF did not improve the stability of lyophilized CFPS reactions to high temperatures or extended storage times. It is possible that the reconstitution of SF-CFPS activity after drying could be improved if the protein loading ratio or drying process is optimized to maintain solubility of the SF.

Beyond the impact of SF on CFPS performance, there is significant potential for CFPS embedded in hydrogels to provide novel functionality to materials. For example, genetic sensors could effect a color change, control material properties such as stiffness, form a gel by producing a crosslinking enzyme, or degrade a gel by expressing an appropriate protease. Importantly for some applications, protein-based hydrogels embedded with CFPS could provide a genetically functionalized material that is fully biodegradable and requires no GMOs. Disappointingly, the techniques for creation of SF hydrogels by enzymatic crosslinking were inhibited in the presence of CFPS components. Nonetheless, we did observe the intriguing result that crosslinking reagents do not alter CFPS activity, but do result in aggregation as observed by fluorescence microscopy. Aggregation could result from partial crosslinking of the SF or non-specific activity on components of the CFPS mixture. This method to cause aggregation of protein components of the CFPS reaction without activity loss could have applications for engineering localization in cell-free systems. To achieve a silk hydrogel functionalized with CFPS, improvements to the system could include optimization of redox conditions in the reaction or further screening of crosslinking enzyme variants that, for example, have a more specific crosslinking mechanism that performs better in the complex lysate environment.

While there are drawbacks to the use of SF fibroin to form SF-CFPS hydrogels, alternate protein-based gel matrices like elastin, collagen, or *de novo* designed peptides may also be used to embed cell lysates. There are also a plethora of synthetic hydrogel materials that could be suitable for CFPS applications. These materials have differing gelation mechanisms, and could result in improved preservation and material properties. Exploration of this space is an important subject of future work.

## Acknowledgements

We thank our ARAP collaborators at AFRL and NRL for supplying tyrosinase plasmid, SF samples, and expertise. We thank Stephanie Cole for reagents and helpful conversations on cell-free systems, and Scott Walper for providing purified sfGFP. We also recognize our funding sources: OSD SBME ARAP and the US Army CCDC CBC BEAMS program. This work was performed while Marilyn Lee held an NRC Research Associateship award at US Army CCDC CBC.

## Abbreviations

CFPS: Cell-free Protein Synthesis
SF: Silk Fibroin
HRP: horse radish peroxidase
sfGFP: superfolder green fluorescent protein
SEM: Scanning Electron Microscopy

