## Supplementary Information for "Silk Fibroin as an Additive for Cell-Free Protein Synthesis"

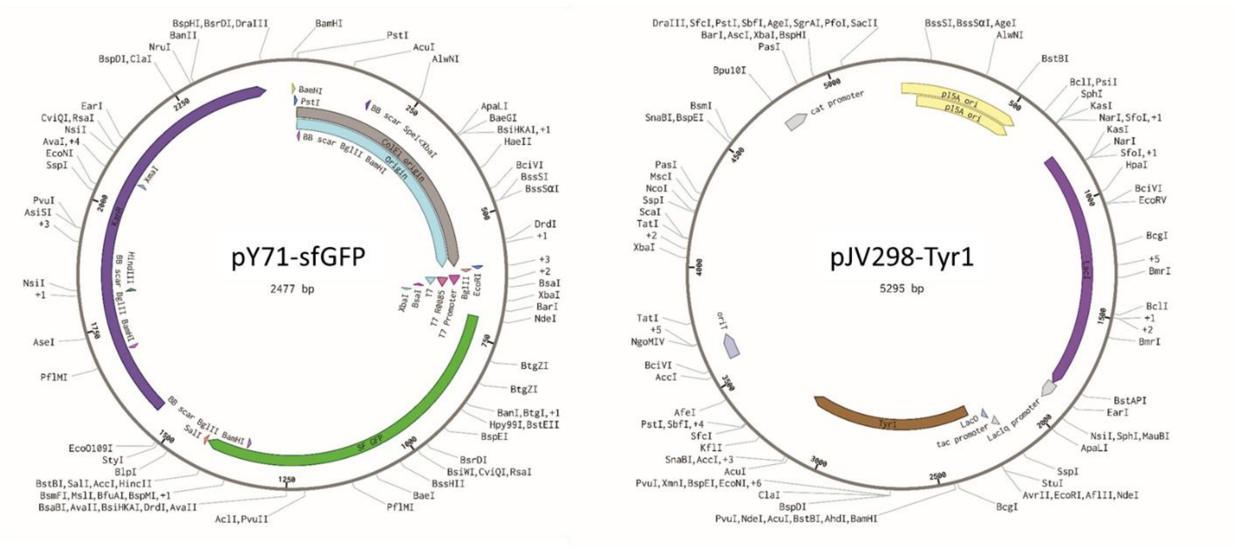


Figure S1. Plasmid maps of pY71-sfGFP (left) and pJV298-Tyr1 (right)

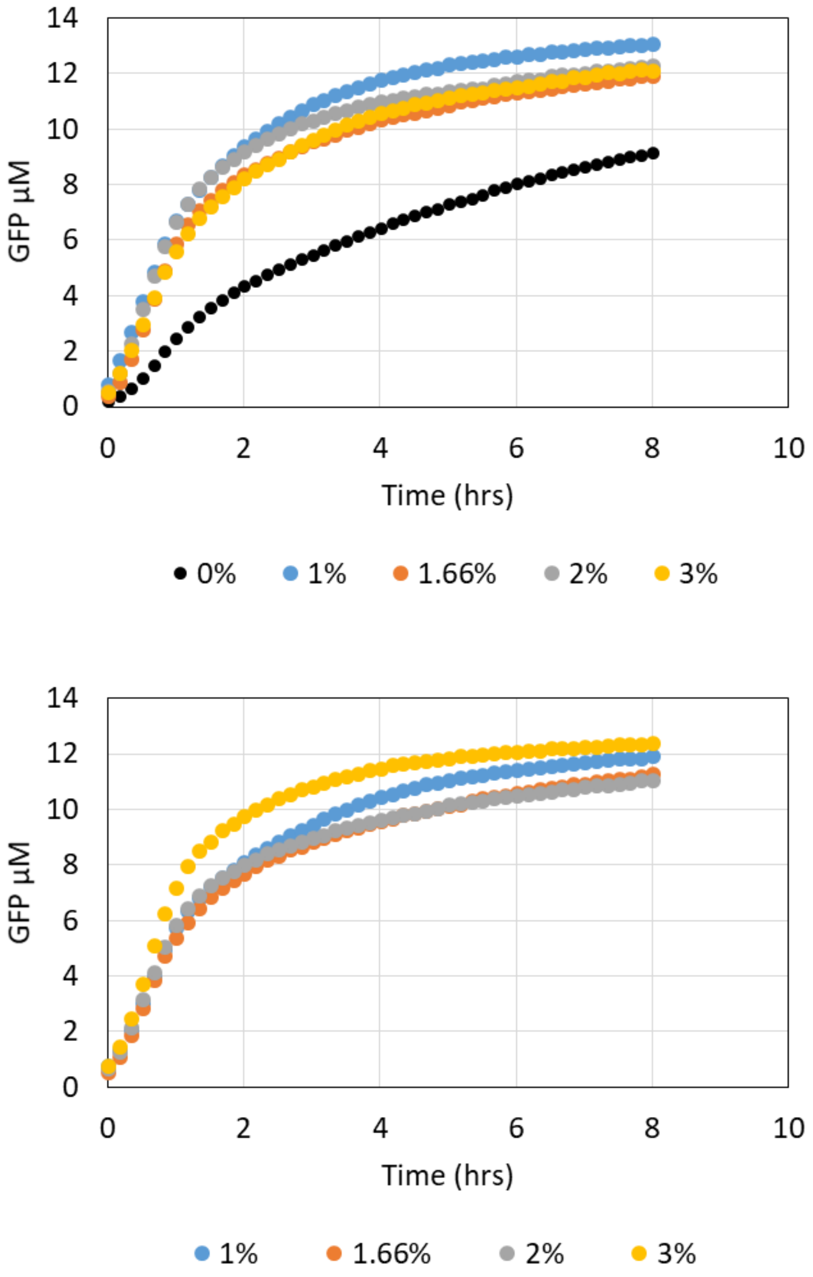


Figure S2. Time course of CFPS reaction producing GFP in the presence of various concentrations of SF with $H_2O_2$ and HRP crosslinking (bottom) or without (top). Each point represents the mean (n≥ 4). Error bars are not included for clarity, refer to Figure 1 of the main text for endpoint and kinetics summary with error bars.

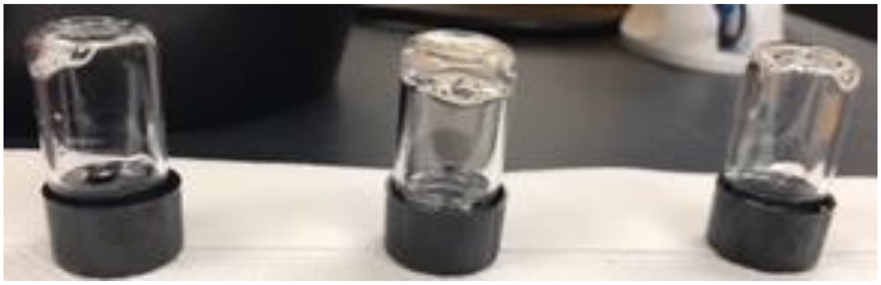

CFPS +SF

SF

Lysate only +SF

Figure S3. Vial inversion test for hydrogel formation. All samples contain 1% SF solution, HRP, $H_2O_2$, and water. In addition the left sample contains all CFPS components. The middle sample has no additions. The right sample included added *E. coli* lysate. Only the middle sample with no CFPS ingredients included exhibits hydrogel formation; the other two liquid samples flow into the cap when the vial is inverted.

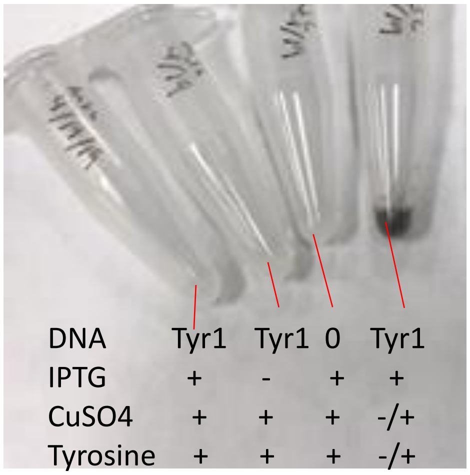


Figure S4. CFPS reactions in 1.5 mL tubes expressing Tyr1 (Tyrosinase, *Bacillus megaterium*). Tyr1 production via CFPS is initiated by addition of purified pJV298-Tyr1 plasmid DNA encoding Tyr1 under the regulatory control of a tac promoter with a lac operator. IPTG is also added to induce expression. The concentrated tyrosine solution was found to inhibit CFPS production of GFP. Therefore, to test Tyr1 activity tyrosine and $CuSO_4$ solutions were added after 2 hours incubation of the protein synthesis reaction to initiate melanin formation (indicated by a -/+ symbol). The optimal conditions yielded dark melanin pigment indicating Tyr1 activity. Tube labeled ‘0’ indicates a control with no DNA.

Detailed Lysate Preparation Protocol

**Media and Buffer Preparation**

To make 2L worth of cells for sonication:

Prepare 2.25L of 2xYPTG media. The final volume will be adjusted after the addition of glucose post-autoclaving.

Reduce final volume of media by 1/10 of the volume to account for the addition of glucose:

100mL glucose to 900mL media for 1L
50mL glucose to 450mL media for 0.5L
10mL glucose to 90mL media for 0.1L

**2xYPTG media**

| Component | For a final volume of 1L | For a final volume of 2.5L |
|---|---|---|
| Tryptone | 16g | 40g |
| Yeast Extract | 10g | 25g |
| NaCl | 5g | 12.5g |
| $K_2HPO_4$ Potassium Phosphate Dibasic | 7g | 17.5g |
| $KH_2PO_4$ Potassium Phosphate Monobasic | 3g | 7.5g |
| | Suspend in 0.9 L total V to leave room for glucose | Suspend in 2.25 L total V to leave room for glucose |

Adjust pH to 7.2 with 5 N KOH.

Add 90mL 2xYPT (above mixture pre-glucose addition) to a 500mL baffled flask, and add 450mL each to a total of four 2L baffled flasks. Add the remainder of the media to a bottle.

Make a solution of 54g of glucose in 300mL ultra-pure water (0.18g/mL glucose). A final concentration of 18g/L of glucose will be added to the media. Autoclave or filter sterilize prior to adding to 2xYPTG media.

**S30a Buffer**

To make 1L:

| Component | Stock Concentration | Final Concentration | Amount for 1L |
|---|---|---|---|
| Tris-OAc, pH 8.2 @ RT | 1M | 10mM | 10mL |
| $Mg(OAc)_2$ | 1.4M | 14mM | 10mL |
| KOAc | 6M | 60mM | 10mL |

**Make Mg(OAc)2 fresh since it is contaminated easily

Add ultra-pure water to a final volume of 1L. Autoclave and store at 4°C.

Add DTT just before use. Guidelines for DTT addition are found below.

**Cell Culture**

Pre-chill the floor centrifuge and the benchtop centrifuge to 4°C.

Add 10mL of sterile 0.18g/mL glucose solution to 90mL of autoclaved 2xYPTG media in the 500mL baffled flask.

Add a single aliquot (50uL) of Invitrogen One Shot BL21 Star (DE3) chemically competent E. coli to the 500mL flask. Incubate for 16 hours at 240rpm and 37°C.

After 16 hours, add 50mL of the 0.18g/mL glucose solution to each of the four 2L baffled flasks containing 450mL 2xYPTG media.

Next, use the 100mL overnight culture to seed the four 2L flasks. Add 5mL of the overnight culture to each 2L flask containing 500mL 2xYPTG. The starting OD600 should be around 0.1.

Place the four flasks in a 37°C incubator at 240rpm and periodically check the OD600 until it reaches a value of 3. If the starting OD600 is roughly 0.1, it will take approximately 3 hours 15 minutes.

> The doubling time was calculated to be approximately 33 minutes with an initial lag time of approximately 45 minutes.

After reaching an OD600 of 3.0, centrifuge cells at 5,000xg at 4 °C for 15 min. Four centrifuge bottles are used to each contain 500mL of cells.

Make a 1M DTT solution.

> 154.253g/mol
> Add 0.617g to 4 mL of ultra-pure water (concentration is 0.154g/mL).

Aliquot 0.5L of S30a buffer into a new bottle, and add 1mL of 1M stock for a final concentration of 2mM. Keep remainder of the 1M DTT solution in -20°C freezer as it will be needed the next day in the sonication process.

**Keep S30a buffer in 4°C or ice bucket for duration of the process.

After the initial spin down of the cells, pour off the supernatant and collect in a waste container for decontamination at the end of the process.

**From this point onward, keep the cells chilled at all times.

*Wash 1*

Add approximately 30mL of S30a buffer to each centrifuge bottle and vortex until the pellet becomes dislodged from the bottom. At this time, transfer the contents of the centrifuge bottle to a 50mL conical tube. Repeat for the remainder of the centrifuge bottles.

Continue to vortex each 50mL conical tube until the pellet is completely re-suspended. Return tubes to an ice bucket periodically during vortexing to keep cells chilled. Bring all volumes up to the same amount with S30a buffer prior to centrifuging.

In the chilled benchtop centrifuge, centrifuge the four 50 mL conical tubes at 4,800xg at 4 °C for 10 min.

After centrifuging, pour off the supernatant and collect in a separate waste bottle from the media waste. This waste will need to be turned in due to the DTT.

*Washes 2, 3, and 4*

Add S30a buffer to each conical tube to the 40mL line. Vortex each tube until the pellets are completely re-suspended. Centrifuge again at 4,800xg at 4 °C for 10 min. Pour off the S30a waste, and repeat the washing two more times until 4 total washes are completed.

After the fourth wash, pour off the S30a buffer. Weigh each conical tube with the cap and record the weight of each wet cell pellet. Each empty 50mL conical tube weighs an average of 12.8g. Subtract this value from the weight of the conical tubes to get the net wet cell weight for each pellet.

Each cell pellet usually weighs 3.5-4g (per 500mL of original culture).

At this point, the pellets can be flash frozen in liquid nitrogen and stored at -80 °C until sonication.

**Sonication**

Pre-chill the Eppendorf microtube centrifuge to 4°C prior to starting.

Remove frozen cell pellets from -80 °C freezer and allow them to thaw in an ice bucket.

Add 25mL of S30a buffer into a new bottle and add 50uL of the 1M DTT stock from the previous day that was stored in the -20°C freezer.

Add 1 mL of chilled Buffer S30a per 1 g of cell mass and put the tube in ice bucket for 3-5 min. Vortex to completely re-suspend the pellets.

Once re-suspended, aliquot 1-1.5mL of cell slurry to 1.5mL Eppendorf microcentrifuge tubes. The amount aliquoted will determine the method of sonication.

Choose the sonication method below based on the size aliquots of cell slurry.

| Sonicator Model | Aliquot | Amplitude | On Duty Time | Off Duty Time | Total Energy |
|---|---|---|---|---|---|
| Q125 | 1mL | 25% | 10sec | 10sec | 540J |
| Q500 | 1.5mL | 20% | 40sec | 59sec | 540J |

Once the sonication method is chosen, don hearing protection and begin the sonication process. Lower the sonication probe about halfway into the tube and hit the “pulse” button keeping the probe from touching the walls of the tube. Allow sonication to continue until the correct amount of energy is reached. At that time, press “stop” to end sonication.

After sonicating each tube, add either 3uL (for 1mL lysate) or 4.5uL (for 1.5mL lysate) of 1M DTT per tube and invert to mix. Keep tubes on ice throughout the sonication process.

Once all tubes are sonicated, clean the probe with 70% ethanol and water.

Spin the microcentrifuge tubes at 12,000xg for 10 mins at 4°C.

Remove the supernatant from each tube, remaining cautious to not disturb the pellets. Pool the supernatant in a 50mL conical tube on ice.

2L of cell culture should yield approximately 10mL of lysate.

After pooling all of the supernatant, invert the tube several times to mix. Aliquot in 100uL increments into 1.5mL Eppendorf tubes and immediately flash freeze in liquid nitrogen.

Store frozen aliquots in the -80°C freezer until needed.

Assay for activity using the sfGFP plasmid and compare activity to previous batches.
